# A sound-sensitive source of alpha oscillations in human non-primary auditory cortex

**DOI:** 10.1101/590323

**Authors:** Alexander J. Billig, Björn Herrmann, Ariane E. Rhone, Phillip E. Gander, Kirill V. Nourski, Beau F. Snoad, Christopher K. Kovach, Hiroto Kawasaki, Matthew A. Howard, Ingrid S. Johnsrude

**Author notes:** Alexander J. Billig and Björn Herrmann contributed equally to this work. Corresponding Author: Alexander J. Billig, UCL Ear Institute, 332 Gray’s Inn Road, London, WC1X 8EE, United Kingdom.

## Abstract

The functional organization of human auditory cortex can be probed by characterizing responses to various classes of sound at different anatomical locations. Along with histological studies this approach has revealed a primary field in posteromedial Heschl’s gyrus (HG) with pronounced induced high-frequency (70-150 Hz) activity and short-latency responses that phase-lock to rapid transient sounds. Low-frequency neural oscillations are also relevant to stimulus processing and information flow, however their distribution within auditory cortex has not been established. Alpha activity (7-14 Hz) in particular has been associated with processes that may differentially engage earlier versus later levels of the cortical hierarchy, including functional inhibition and the communication of sensory predictions. These theories derive largely from the study of occipitoparietal sources readily detectable in scalp electroencephalography. To characterize the anatomical basis and functional significance of less accessible temporal-lobe alpha activity we analyzed responses to sentences in seven human adults (four female) with epilepsy who had been implanted with electrodes in superior temporal cortex. In contrast to primary cortex in posteromedial HG, a non-primary field in anterolateral HG was characterized by high spontaneous alpha activity that was strongly suppressed during auditory stimulation. Alpha-power suppression decreased with distance from anterolateral HG throughout superior temporal cortex, and was more pronounced for clear compared to degraded speech. This suppression could not be accounted for solely by a change in the slope of the power spectrum. The differential manifestation and stimulus-sensitivity of alpha oscillations across auditory fields should be accounted for in theories of their generation and function.

**Significance Statement:** To understand how auditory cortex is organized in support of perception, we recorded from patients implanted with electrodes for clinical reasons. This allowed measurement of activity in brain regions at different levels of sensory processing. Oscillations in the alpha range (7-14 Hz) have been associated with functions including sensory prediction and inhibition of regions handling irrelevant information, but their distribution within auditory cortex is not known. A key finding was that these oscillations dominated in one particular non-primary field, anterolateral Heschl’s gyrus, and were suppressed when subjects listened to sentences. These results build on our knowledge of the functional organization of auditory cortex and provide anatomical constraints on theories of the generation and function of alpha oscillations.

## Introduction

Human primary auditory cortex occupies the posteromedial portion of Heschl’s gyrus and can be distinguished from neighbouring non-primary fields based on architectonic and electrophysiological features (Hackett et al., 2001; Clarke and Morosan, 2012; Nourski and Howard, 2015). Responses to sound in primary auditory cortex phase-lock to rates beyond 100 Hz and include characteristic short-latency evoked components (Liégeois-Chauvel et al., 1991; Howard et al., 2000; Brugge et al., 2008, 2009). They also contain pronounced induced power in the high gamma range (70-150 Hz; Steinschneider et al., 2008; Brugge et al., 2009), which is considered a proxy for multi-unit spiking activity (Crone et al., 2001; Mukamel et al., 2005; Manning et al., 2009).

The anatomical distribution and stimulus-sensitivity of ongoing low-frequency oscillations in auditory cortex have been less well characterized, but there are several reasons to expect the strength of alpha-band (broadly defined as 7-14 Hz) activity in particular also to vary across primary and non-primary auditory cortex. First, cortical high-gamma and alpha activity often exhibit an antagonistic relationship (Crone et al., 1998; Mukamel et al., 2005; Ramot et al., 2012). For example, alpha power in visual cortex is suppressed as high gamma activity increases upon afferent stimulation (Hoogenboom et al., 2006; Bauer et al., 2014). Second, primary and non-primary auditory fields have distinct architectonic and connectivity profiles (Clarke and Morosan, 2012), which may give rise to intrinsic dynamics at different timescales (Başar and Güntekin, 2008; Honey et al., 2013; Murray et al., 2014). Third, alpha oscillations have been associated with a range of neural operations and processes that may differentially engage lower versus higher levels of the cortical hierarchy (Clayton et al., 2018). These include functional inhibition (Klimesch et al., 2007; Jensen and Mazaheri, 2010), inter-areal synchronization (Palva et al., 2010), and the communication of sensory predictions (Bauer et al., 2014; Sedley et al., 2016; Auksztulewicz et al., 2017; Chao et al., 2018).

Most of what we know of human alpha oscillations derives from study of the dominant occipital and parietal sources detectable in scalp electroencephalography (EEG), particularly when the eyes are closed or during manipulations of visuo-spatial attention (Berger, 1931; Clayton et al., 2018). However, ongoing oscillations at the lower end of the alpha frequency range (7-10 Hz) have also been recorded from superior temporal cortex. These show power decreases during auditory stimulation (Niedermeyer, 1990; Tiihonen et al., 1991; Lehtelä et al., 1997; Weisz et al., 2011; Fontolan et al., 2014), the extent of which is sensitive to the nature of the stimulus, for example with clear speech eliciting a larger power suppression compared to noisy speech (Obleser and Weisz, 2012; de Pesters et al., 2016). There are indications that the magnitude of ongoing alpha oscillations (Frauscher et al., 2018) and of their suppression during stimulation (Fontolan et al., 2014) is lower on Heschl’s gyrus than elsewhere in the temporal lobe. However, recording sites in the relevant studies were not localized to primary versus non-primary auditory cortex based on known electrophysiological response properties.

In order to more precisely characterize the spatial distribution and time-course of low-frequency oscillations, including alpha, in human auditory cortex, we recorded from eight hemispheres in seven adult patients implanted with electrodes along the length of Heschl’s gyrus and elsewhere on the superior temporal plane and lateral superior temporal gyrus, during clinical monitoring for epilepsy. This coverage allowed analysis of local field potentials in both primary and non-primary auditory fields, defined both anatomically and based on responses to click trains, while subjects listened to clear and degraded speech. Our objective was not to directly test particular accounts of alpha generation or function, but rather to provide anatomical specificity that may constrain the development of these theories, particularly as they relate to auditory processing.

## Materials and Methods

### Subjects

Subjects for the speech experiment were seven individuals (four females; median age: 33 years; age range 22-56 years) undergoing intracranial monitoring for diagnosis and treatment of medically intractable epilepsy. Recordings were made in an electromagnetically shielded hospital room at the Epilepsy Monitoring Unit at the University of Iowa Hospitals and Clinics. Three individuals had electrode placement in the left hemisphere only (subjects L357, L403, L442), three in the right hemisphere only (subjects R369, R399, R429) and one bilaterally (subject B335). Further details of electrode placement are provided in the “Data recording” section and in Figure 1. All subjects were right-handed with left-hemisphere language dominance as determined by preimplantation Wada testing. Three subjects (B335, R399, L442) had normal hearing (pure tone thresholds below 20 dB hearing level for frequencies between 250 and 4000 Hz) and four (L357, R369, L403, R429) had mild hearing loss at isolated frequencies (there was no systematic difference in results between these two sub-groups). Vision was self-reported as normal or corrected to normal; participants who required glasses wore them during the task. All participants were native speakers of English. Subject details, including demographic data, seizure focus, details of hearing loss, and the number of contacts in each studied field, are provided in Table 1.

**Table 1.** Demographics, hearing, seizure focus, and electrode coverage

| Subject | Age (years) | Sex | Hearing loss in dB HL (hearing level) with frequency in kHz and ear | Seizure focus | Contacts by regions and hemisphere |  |  |  |  |  |
| --- | --- | --- | --- | --- | --- | --- | --- | --- | --- | --- |
|  |  |  |  |  | pmHG |  | alHG |  | Other STP and STG |  |
|  |  |  |  |  | L | R | L | R | L | R |
| B335<sup>a</sup> | 33 | M | None | Bilateral medial temporal | 2 | 4 | 4 | 6 | 10 | 8 |
| L357 | 36 | M | 30 dB, 4 kHz, L | L posterior hippocampus | 6 | - | 5 | - | 9 | - |
| R369 | 30 | M | 30 dB, 4 kHz, L | R medial temporal | - | 8 | - | 5 | - | 31 |
| R399 | 22 | F | None | R temporal | - | 3 | - | 3 | - | 28 |
| L403 | 56 | F | 30 dB, 0.5 kHz, L; 25 dB, 4 kHz, L | L posterior hippocampus | 8 | - | 4 | - | 30 | - |
| R429 | 32 | F | 25 dB, 2 kHz, L | R medial temporal | - | 2 | - | 3 | - | 13 |
| L442 | 36 | F | None | L temporal | 6 | - | 6 | - | 33 | - |
| **Total** |  |  |  |  | 39 |  | 36 |  | 162 |  |

<sup>a</sup> left and right contacts are analyzed and presented separately as L335 and R335; pmHG – posteromedial Heschl's gyrus; alHG – anterolateral Heschl's gyrus; STP – superior temporal plane; STG – superior temporal gyrus; M – male; F – female; L – left; R – right

**Figure 1.**
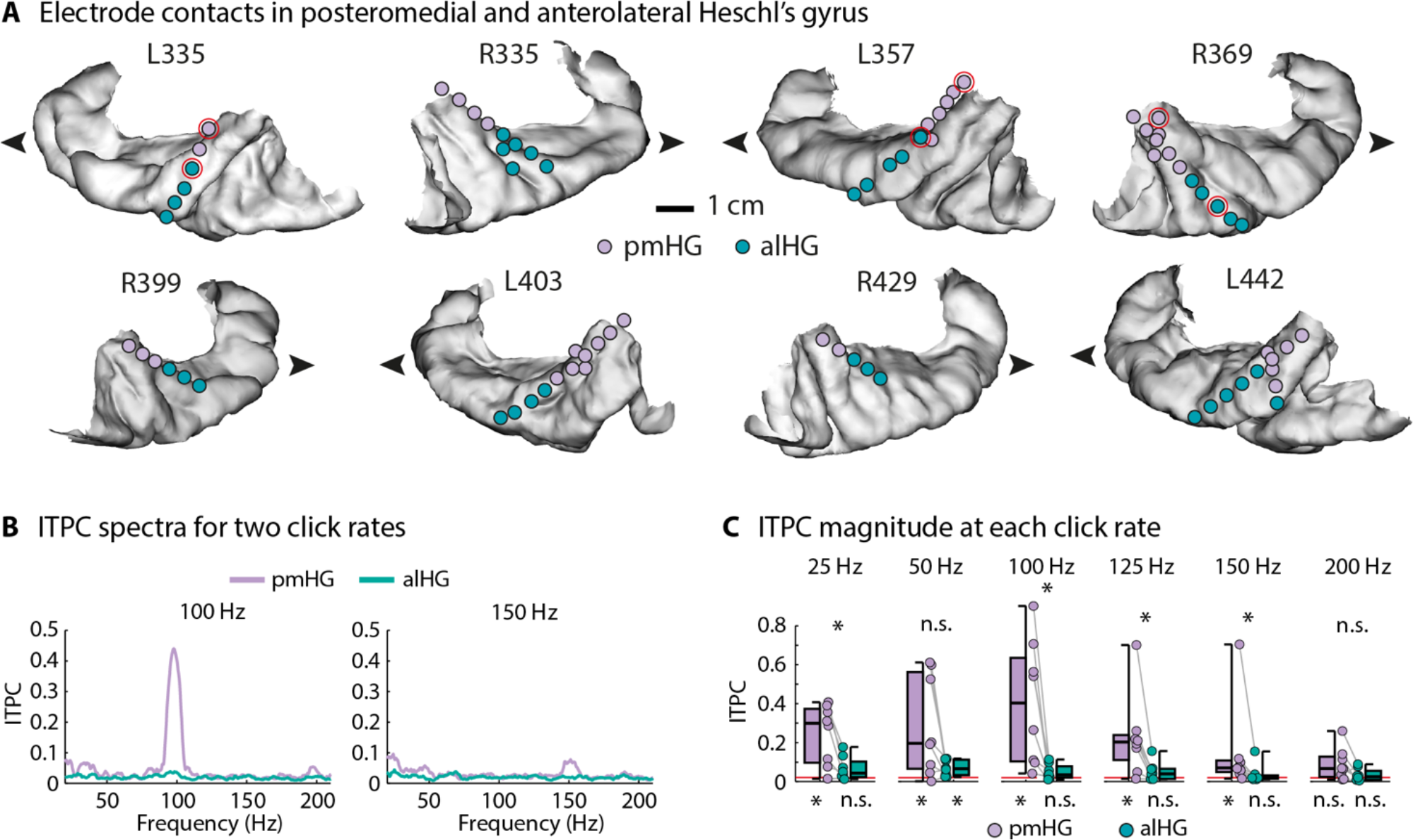
Anatomical and functional separation of primary and non-primary auditory cortex on Heschl’s gyrus. **A:** Recording contacts functionally identified as primary auditory cortex in posteromedial Heschl’s gyrus (pmHG) are in purple and those functionally identified as non-primary auditory cortex in anterolateral Heschl’s gyrus (alHG) are in green. Pairs of contacts circled in red for L335, L357, and R369 are those for which sample trials are plotted in Figure 2. Contacts outside of Heschl’s gyrus are not shown. The lateral surface of the superior temporal gyrus forms the lower bound of each image. The arrowhead indicates the anterior direction. The size of symbols depicting recording contacts has been increased for clarity. **B:** Inter-trial phase coherence (ITPC) for two example click rates (100 Hz and 150 Hz; median across hemispheres). **C:** ITPC at the neural frequencies corresponding to the click repetition rate, for the six different click rates. Data points reflect individual hemispheres, and box and whiskers indicate the semi-interquartile and full range. The horizontal red line close to zero indicates chance level. Asterisks below indicate a significant ITPC difference from chance level, and asterisks above reflect a significant difference between pmHG and alHG (*p* ≤ 0.05, FDR-corrected, n.s.: not significant). These differences arise by definition: significant phase-locking at high stimulus rates was one of the criteria used to assign recording sites to areas.

Research protocols were approved by the University of Iowa Institutional Review Board, and subjects signed informed consent documents prior to any recordings. Research did not interfere with acquisition of clinical data, and subjects could withdraw from research at any time without consequence for their clinical monitoring or treatment. Subjects initially remained on their antiepileptic medications but these were typically decreased in dosage during the monitoring period at the direction of neurologists until sufficient seizure activity had been recorded for localization, at which point antiepileptic medications were resumed. No research occurred within three hours following seizure activity. Electrode contacts at seizure foci were excluded from all analyses.

### Stimuli

Stimuli for determining primary versus non-primary sites were 160-ms long click trains, presented at rates of 25, 50, 100, 125, 150 and 200 Hz. 50 click trains were presented at each rate, in random order and with intervals between onsets of successive click trains drawn from a normal distribution with mean 2000 ms and standard deviation 10 ms. Stimuli for the main experiment were clear and noise-vocoded versions of English sentences previously used by Wild et al. (2012). They were recorded by a female native speaker of North American English in a soundproof booth using an AKG C1000S microphone with 16-bit sampling at 44.1 kHz using an RME Fireface 400 audio interface. Three-band noise-vocoding was performed as described by Shannon et al. (1995) using a custom-designed vocoder implemented in Matlab. In detail, items were filtered into three contiguous frequency bands (50-558 Hz, 558-2264 Hz, 2264-8000 Hz, selected to have equal spacing along the basilar membrane), using FIR Hann band-pass filters with an 801-sample window length. The amplitude envelope in each frequency band was extracted using full-wave rectification followed by low-pass filtering at 30 Hz with a fourth-order Butterworth filter. These envelopes were applied to band-pass filtered noise in the same frequency ranges, the results of which processing were summed to produce the noise-vocoded sentence. Clear speech remained unprocessed, containing all frequencies up to 22.05 kHz.

### Procedure

Subjects took part while sitting upright in their hospital bed. Sounds were presented diotically at a comfortable listening level via insert earphones (ER4B, Etymotic Research) integrated into custom-fit earmolds. For the presentation of click trains, subjects relaxed and performed no task. For the main experiment, subjects fixated on the centre of a screen (ViewSonic VX922 or Dell 1707FPc) positioned approximately 60 cm in front of them and made responses on a computer keyboard. Each trial consisted of an initial auditory sentence presentation (either clear or degraded; duration 1223-4703 ms), a delay (1250-1500 ms), a written sentence (2500-2750 ms), a further delay (1250-1500 ms), a repeat presentation of the auditory sentence, and a further delay (2300-2800 ms). During the delay periods a fixation cross was displayed in the centre of the screen. The written sentence either matched, or was completely different to, the content of the spoken sentence. To facilitate attention to the auditory stimuli, subjects were told that they would be asked about the content of the spoken sentences at the end of the experiment. To encourage attention to the text, subjects were asked to report occasional capital letters that occurred in 9% of trials (excluded from analysis) using the hand ipsilateral to the hemisphere with the most electrode contacts. Stimulus presentation and response collection was controlled by Presentation (Neurobehavioral Systems, Inc., Berkeley, CA, USA) running on a Windows PC. The speech experiment lasted between 30 and 45 minutes, depending on the number of trials completed (range 120-136) and whether the subject opted to take a break. The manipulation of the visual cue was included to answer a separate research question concerning the neural basis of perceptual pop-out of degraded speech; related results including performance in the behavioral tasks will be reported elsewhere.

### Data recording

Electrophysiological activity was recorded using depth and subdural electrodes (Ad-Tech Medical, Racine, WI). Subdural grids with coverage including superior temporal gyrus consisted of platinum-iridium discs (2.3 mm diameter, 5-10 mm inter-contact distance) embedded in a silicon membrane. Depth arrays of 8-12 recording contacts spaced at distances of 5 mm were implanted stereotactically, targeting Heschl’s gyrus. In some patients, additional arrays targeting the insula provided coverage of other sites in the superior temporal plane (Nagahama et al., 2018). A subgaleal contact served as a reference. Electrode placement was determined solely on the basis of clinical requirements, as determined by the neurosurgery and epileptology team (Nourski and Howard, 2015). Data acquisition was via a TDT RZ2 real-time processor (Tucker-Davis Technologies, Alachua, FL, USA) for subjects B335 and L357, and via a Neuralynx Atlas System (Neuralynx Inc., Bozeman, MT, USA) for the remaining subjects. Data were amplified, filtered (TDT: 0.7-800 Hz bandpass, 12 dB/octave rolloff; Neuralynx: 0.1-500 Hz, 5 dB/octave rolloff, with the exception of subject R369 for whom filtering was at 0.8-800 Hz), and digitized with a sampling rate of 2034.5 Hz (TDT) or 2000 Hz (Neuralynx).

Anatomical reconstruction of implanted electrode location, mapping to a standardized coordinate space, and assignment to regions of interest were performed using FreeSurfer image analysis suite (version 5.3; Martinos Center for Biomedical Imaging, Harvard, MA) and custom software, as described previously (Nourski et al., 2014). In brief, whole-brain high-resolution T1-weighted structural magnetic resonance imaging (MRI) scans (resolution and slice thickness ≤1.0 mm) were obtained from each subject before electrode implantation. After implantation, subjects underwent MRI and thin-slice volumetric computerized tomography (CT) (resolution and slice thickness ≤1.0 mm) scanning. Electrode locations were initially extracted from post-implantation MRI and CT scans, then projected onto preoperative MRI scans using non-linear three-dimensional thin-plate spline morphing, aided by intraoperative photographs. Where required, standard Montreal Neurological Institute (MNI) coordinates were found for each contact using linear coregistration to the ICBM152 atlas, as implemented in FMRIB Software library (Version 5.0; FMRIB Analysis Group, Oxford, UK).

### Data analysis

Offline data analysis was carried out in Matlab Software (MathWorks Inc.; version 2012b) using the Fieldtrip Toolbox (http://fieldtrip.fcdonders.nl/; v20131231; Oostenveld et al., 2011) and custom Matlab scripts. Power line noise was removed using a filter based on the demodulated band transform (Kovach and Gander, 2016). Data were then downsampled to 1000 Hz, and recordings divided into epochs from –1500 to 6300 ms relative to sentence onset.

Responses to click trains were investigated using inter-trial phase coherence (ITPC; Lachaux et al., 1999), which indexes of the strength with which neural activity synchronizes (phase locks) with temporally regular patterns in acoustic stimulation (e.g. Herrmann and Johnsrude, 2018). To this end, a fast Fourier transform (including a Hann window taper and zero-padding) was calculated for frequencies ranging from 20–210 Hz using the data in the 0–0.2 s time window (i.e., the click-train duration). The resulting complex numbers were normalized by dividing each by its magnitude. ITPC was then calculated as the absolute value of the mean normalized complex number across trials. ITPC can take on values between 0 (no coherence) and 1 (maximum coherence). ITPC was calculated separately for each trial, contact, and repetition rate (50, 100, 125, 150, 200 Hz). We also calculated ITPC based on surrogate data in order to compare the empirically observed ITPC values against ITPC chance levels (Stam, 2005). In detail, time series data (0–0.2 s after stimulus onset) were converted to the frequency domain using an FFT; the phase of each frequency component was randomized, and the data were subsequently converted back to the time domain. ITPC was then calculated for the surrogate time series. Calculation of surrogate data and ITPC was repeated 100 times, and ITPC values were averaged subsequently, leading to a chance-level ITPC.

For the main experiment, time-frequency analysis of oscillatory activity was conducted separately for low-frequency (2-30 Hz) and high-frequency activity (40–180 Hz) activity using Morlet wavelets (Tallon-Baudry et al., 1996; Tallon-Baudry and Bertrand, 1999). In detail, for each sentence and contact, time-frequency representations were calculated for the –0.6–1.2-s time window relative to stimulus onset in 10 ms steps. We limited our analysis time window to 1.2 s in order to ensure that our analysis would be related to ongoing activity during stimulus presentation (1.223 s was the duration of the shortest sentence). For frequencies between 2 and 30 Hz (calculated in steps of 0.2 Hz), wavelet size linearly increased from 3 to 12 cycles as a function of frequency. For frequencies between 40 and 180 Hz (calculated in steps of 1 Hz), a wavelet size of 12 cycles was used uniformly. Power was calculated as the squared magnitude of the complex wavelet coefficients and averaged across trials. Time-frequency power was baseline-corrected by dividing the power at each time point by the mean power in the –0.6 to –0.1 s pre-stimulus time window, taking the logarithm (base 10), and multiplying the result by 10 (separately for each frequency). The result is power in decibel (dB) units, reflecting the signal change from baseline.

### Functional localization

Recording sites were identified as belonging to a primary region of interest in posteromedial Heschl’s gyrus (pmHG) if they showed significant phase-locked responses to 100-Hz click trains, and if averaged click-evoked potentials included short-latency (<20 ms) components (Brugge et al., 2009; Nourski et al., 2016). Other sites along the gyrus that did not demonstrate these properties were deemed to be in non-primary cortex and assigned to the anterolateral Heschl’s gyrus (alHG) region of interest.

### Experimental design and statistical analysis

Non-parametric Wilcoxon signed-rank tests were calculated using the signrank function in Matlab. False discovery rate was used to correct for multiple comparisons (Benjamini and Hochberg, 1995; Genovese et al., 2002), if not indicated otherwise. Effect sizes are reported as *r*_*e*_(*r*_*equivalent*_; Rosenthal and Rubin, 2003), which is equivalent to a Pearson product-moment correlation for two continuous variables, to a point-biserial correlation for one continuous and one dichotomous variable, and to the square root of partial *η*^2^ for ANOVAs.

## Results

### Functional segregation into primary and non-primary auditory cortex

A posteromedial-anterolateral boundary along Heschl’s gyrus between primary and non-primary auditory cortex was identified for each hemisphere (Figure 1A), based on responses to click trains. As per our definition, phase-locking at 100-150 Hz click stimulation rates was significantly greater at posteromedial than at anterolateral contacts (Figure 1B, 1C).

### Alpha-power suppression and high gamma-power enhancement dominate in anterolateral and posteromedial Heschl’s gyrus respectively

Figure 2 shows activity from 1s prior to sentence onset until 1s after sentence onset for two exemplary trials at three pairs of contacts, each consisting of one pmHG and one alHG contact from a single subject and hemisphere. A striking feature is the prominent low frequency pre-stimulus activity in alHG that is reduced following sentence onset. This observation is supported by a group analysis. Figure 3A displays time-frequency power for the low-frequency range (2–30 Hz) for pmHG and alHG, averaged across all recording sites. Motivated by previous research on alpha oscillatory activity in the temporal lobe, we focused on the 7 to 10 Hz frequency band (Tiihonen et al., 1991; Lehtelä et al., 1997).

**Figure 2.**
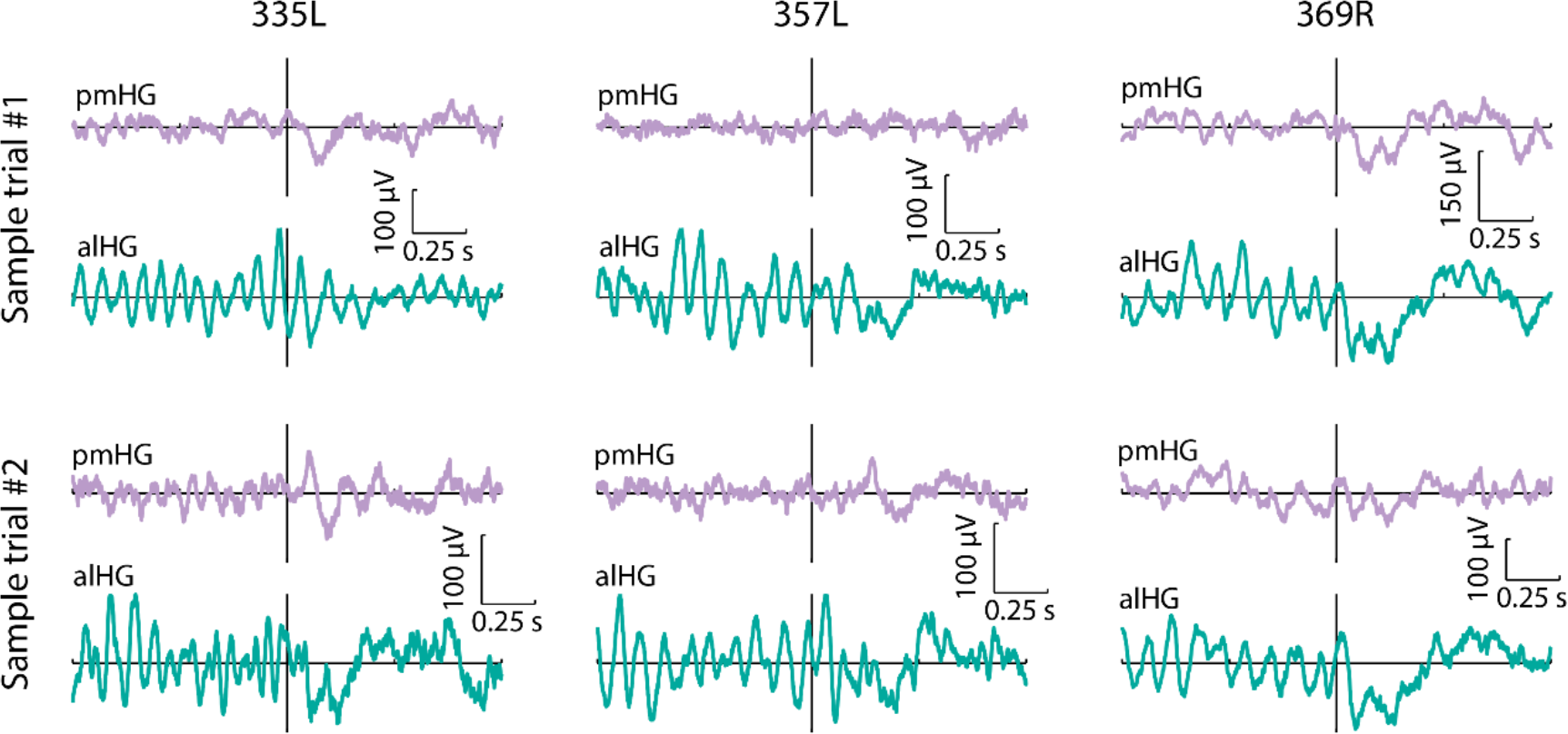
Exemplary single trial activity at pairs of posteromedial and anterolateral Heschl’s gyrus contacts. Raw voltage traces from 1s before sentence onset to 1s after sentence onset for two representative trials as recorded in three hemispheres. For each hemisphere, traces at one pmHG and one alHG contact are plotted; these contacts are outlined in red in Figure 1A. Vertical line indicates stimulus onset.

**Figure 3.**
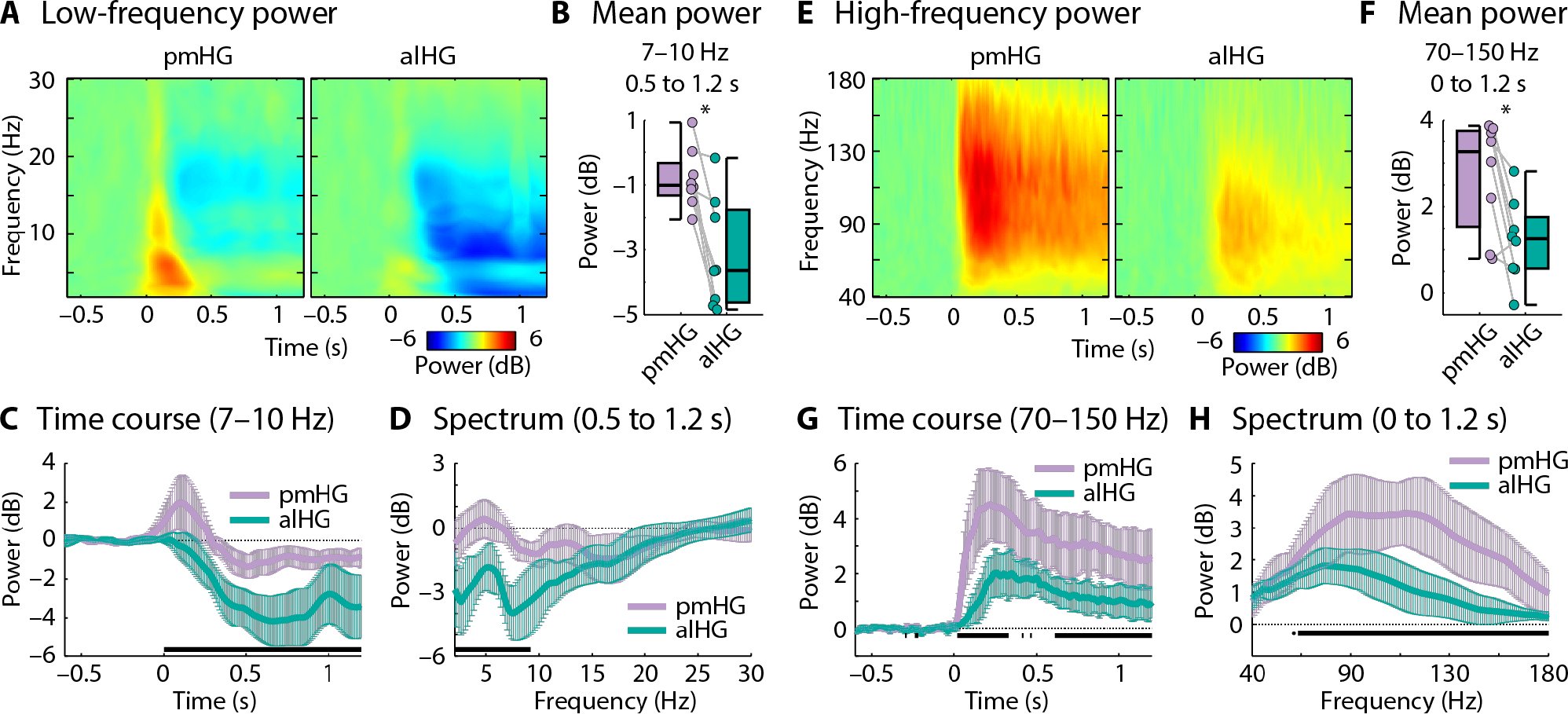
Alpha power and high gamma power changes time-locked to sentence onset. **A:** Spectrogram (time-frequency) representations of power in the 2–30 Hz frequency range across all pmHG (left) and alHG (right) contacts in all hemispheres. **B:** Power in the alpha (7-10 Hz) range. Data points reflect individual hemispheres, and box and whiskers indicate the semi-interquartile and full range (^*^*p* ≤ 0.05). **C:** Alpha power time course (median across hemispheres; error bar reflects the semi-interquartile range). **D:** Low frequency power spectrum (median across hemispheres; error bar reflects the semi-interquartile range). Black line marks significant differences between pmHG and alHG (*p* ≤ 0.05; FDR-corrected). **E-H**: Same as A-D but for the high frequency range. pmHG - posteromedial Heschl’s gyrus; alHG - anterolateral Heschl’s gyrus.

Alpha-power was significantly lower in alHG compared to pmHG (*p* = 0.008, *r*_*e*_ = 0.812; Figure 3B). The time course for the alpha frequency band was extracted and is shown in Figure 3C. Alpha-power suppression after sentence onset (relative to the pre-stimulus period) was significantly stronger in alHG compared to pmHG for the duration of the 0.5-1.2-s analysis window (*p* ≤ 0.05, FDR-corrected, black solid line in Figure 3C). Note that although our analysis focused on 7–10 Hz - motivated by previous work and by the peak in the spectrum (Figure 3D) - the suppression of power in alHG relative to pmHG was also present for lower frequencies. In order to investigate whether the suppression of alpha power persisted for the duration of the sentences, which were of variable length, we also analyzed the data time-locked to sentence offset (baseline-corrected using the 0.1–0.6 s time window post-sentence offset). Alpha power (7–10 Hz; –1.2 to –0.5 s) remained significantly lower in alHG than in pmHG (*p* = 0.023, *r*_*e*_ = 0.737; not plotted) and was below the post-sentence baseline.

The pattern of activity across primary and non-primary cortex in the high gamma range had a different profile. Figures 3E–3H show that the strength of high gamma responses (70-150 Hz) to sentences was greater in pmHG compared to alHG (*p* = 0.023, *r*_*e*_ = 0.737) from shortly after stimulus onset, consistent with previous findings for click trains and single syllables (Brugge et al., 2009; Steinschneider et al., 2014).

### Alpha-power suppression throughout superior temporal cortex decreases with distance from anterolateral Heschl’s gyrus

We wanted to establish whether alpha-power suppression decreases with distance from alHG throughout the superior temporal plane and superior temporal gyrus, which would suggest that the main local source of alpha oscillations is located in alHG. Figure 4A shows alpha power (0.5–1.2 s; 7–10 Hz) relative to the pre-stimulus time window for all superior-temporal-cortex contacts as a function of spatial (Euclidean) distance to the mean coordinate of each subject’s alHG contacts. For each hemisphere, we fitted a linear function to the alpha-power values as a function of spatial distance from this point and tested the estimated linear coefficient against zero. Alpha-power suppression decreased as spatial distance from alHG increased (*p* = 0.023, *r*_*e*_ = 0.737), suggesting that auditory alpha-power suppression originates in alHG.

**Figure 4.**
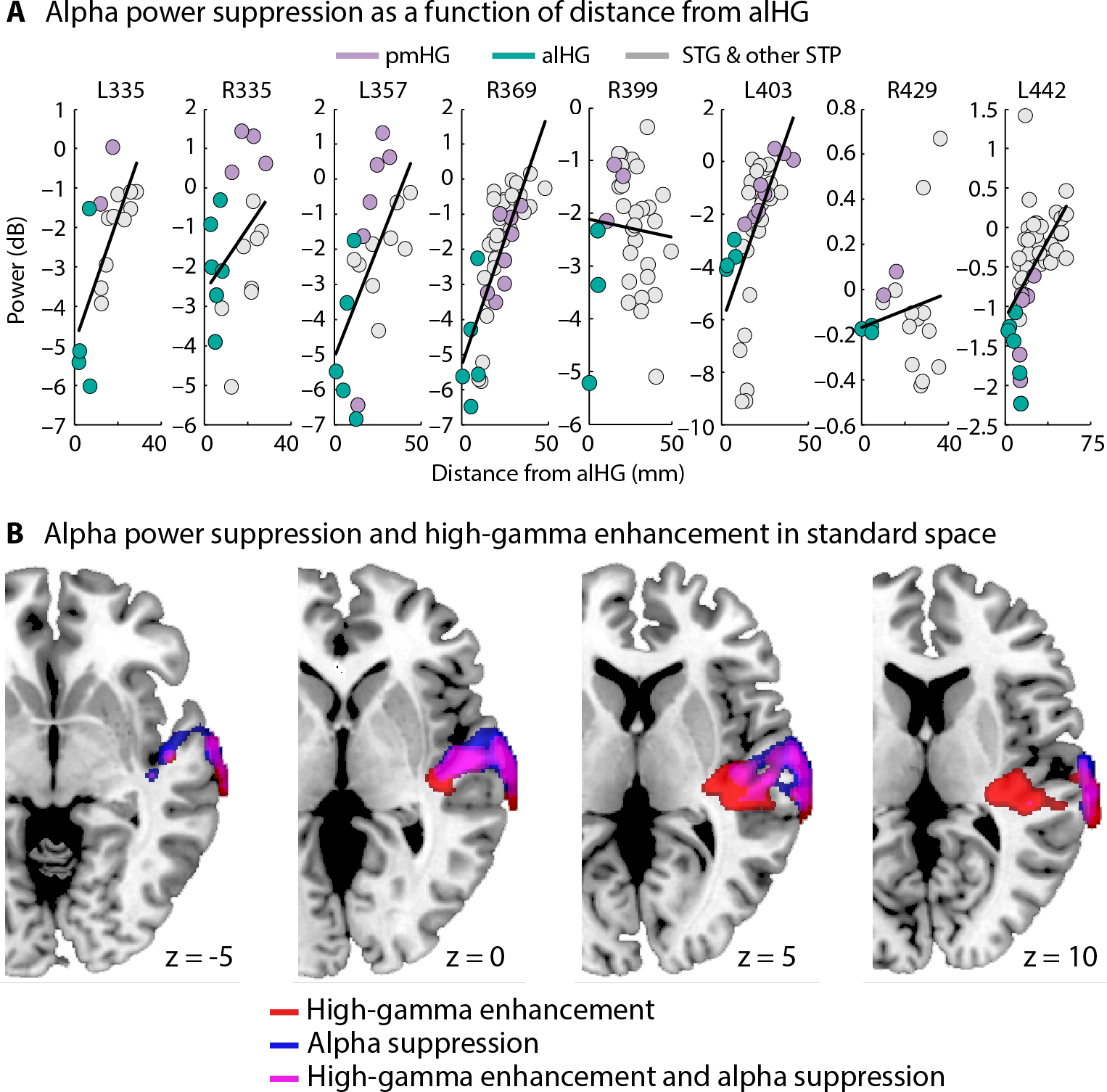
Spatial distribution of alpha-power suppression and high gamma-power enhancement throughout superior temporal cortex. **A:** Baseline-corrected alpha (7-10 Hz; 0.5-1.2 s) power for each contact as a function of distance from the mean coordinate of each subject’s alHG contacts. The black line reflects the best-fitting linear function, estimated separately for each hemisphere. pmHG - posteromedial Heschl’s gyrus; alHG - anterolateral Heschl’s gyrus; STG - superior temporal gyrus; STP - superior temporal plane. **B:** High gamma-power enhancement (70-150 Hz) and alpha-power suppression (7-10 Hz) mapped onto four equally spaced axial slices of a template brain. Data from the left hemisphere were project to the right hemisphere before averaging across subjects. High gamma power is greatest in posterior auditory areas, including on the lateral surface, whereas alpha power is suppressed most strongly in anterior areas.

In order to further visualize activity centres of alpha-power suppression and compare them to centres of high gamma power enhancement, alpha- and high gamma power for each contact were projected onto a template brain (and left-hemisphere contacts were mapped onto the right hemisphere; smoothing 6 mm FWHM; Figure 4B). High gamma-power enhancement (relative to baseline) was most prominent in posteromedial auditory cortex, but also present in the posterior part of the lateral surface of the temporal cortex. Alpha-power suppression (relative to baseline) was strongest in more anterior areas.

### Alpha-power suppression in anterolateral Heschl’s gyrus is greater for clear than for noise-vocoded speech

Previous EEG work suggests that alpha-power suppression is stronger for clear compared to noise-vocoded speech (Obleser and Weisz, 2012). Estimated sources of this effect include extensive parieto-occipital and anterior temporal regions, but the resolution of this localization using scalp recordings is limited. Figure 5 shows that alpha-power suppression in the present data was larger for clear speech compared to noise-vocoded speech in alHG (*p* = 0.039, *r*_*e*_ = 0.692), but not in pmHG (*p* = 0.844, *r*_*e*_ = 0.077), although the stimulus by region interaction was not significant (*p* = 0.148, *r*_*e*_ = 0.523). This indicates that alpha suppression in at least one auditory cortical field is dependent on the spectral quality of the stimulus.

**Figure 5.**
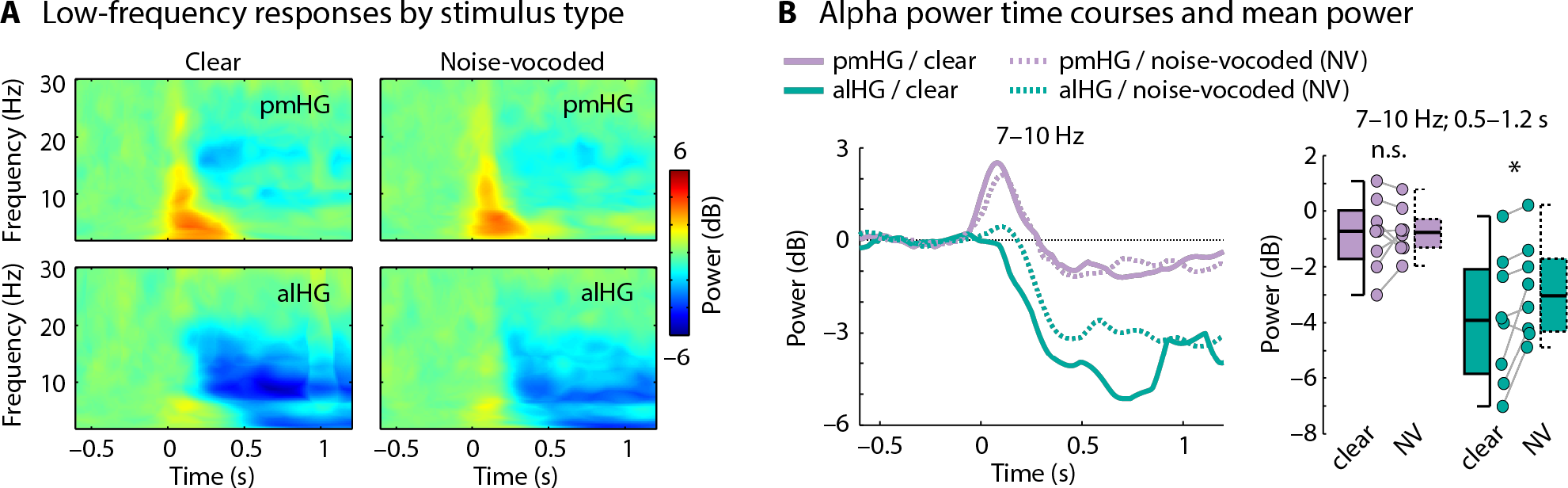
Low-frequency responses for clear speech compared to noise-vocoded speech. **A:** Spectrogram (time-frequency) representations of power in the 2–30 Hz frequency range across all pmHG (top) and alHG (bottom) contacts in all hemispheres for clear (left) and noise-vocoded (right) speech. **B:** Time courses of alpha (7–10 Hz) power. Box plots show alpha power for the 0.5 to 1.2 s time interval. Data points reflect individual hemispheres, and box and whiskers indicate the semi-interquartile and full range. ^*^*p* ≤ 0.05; n.s. - not significant. pmHG - posteromedial Heschl’s gyrus; alHG - anterolateral Heschl’s gyrus.

### Pre-stimulus alpha power is greater in anterolateral than in posteromedial Heschl’s gyrus

Next, we examined the degree to which pre-stimulus power contributes to power differences between pmHG and alHG observed after stimulus onset. Power in the alpha-frequency band (7–10 Hz) was larger in alHG than in pmHG during the pre-stimulus time window (–0.6 to –0.1 s; *p* = 0.008, *r*_*e*_ = 0.812), whereas there was no difference in alpha power between alHG and pmHG for the post-stimulus-onset time window (0.5 to 1.2 s; *p* = 0.742, *r*_*e*_ = 0.128; Figure 6A). The baseline-corrected results presented in Figure 3B demonstrate that the interaction between time period and region was significant.

**Figure 6.**
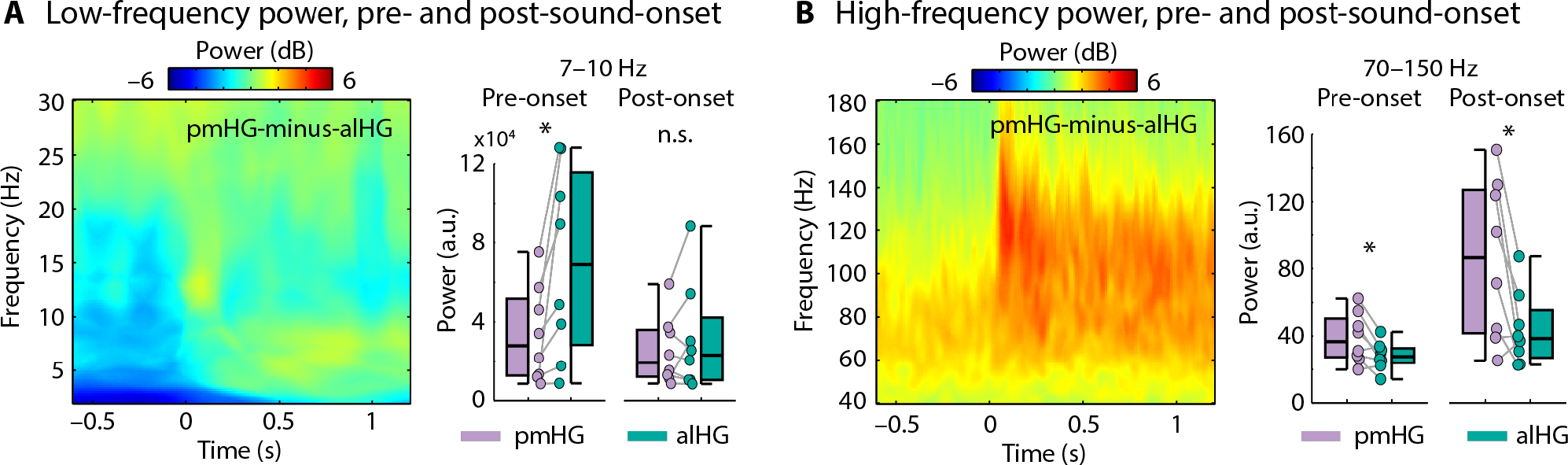
Contrast of posteromedial versus anterolateral Heschl’s gyrus for non-baseline-corrected data. **A:** Spectrogram representation of the difference in low frequency power between pmHG and alHG. Boxplots and individual data points reflect the power in the alpha (7-10 Hz) range for the pre-stimulus time window (–0.6 to –0.1 s) and the post-stimulus-onset time window (0.5 s to 1.2 s) for pmHG and alHG. **B:** Same as panel A, but for the high frequency range, and the post-stimulus-onset time window was 0 s to 1.2 s. pmHG - posteromedial Heschl’s gyrus; alHG - anterolateral Heschl’s gyrus. ^*^*p* ≤ 0.05, n.s. - not significant.

High gamma power (70-150 Hz) was larger in pmHG compared to alHG in both the pre-stimulus (–0.6 to –0.1 s; *p* = 0.039, *r*_*e*_ = 0.692) and in the post-stimulus-onset (0.5–1.2 s; *p* = 0.039, *r*_*e*_ = 0.692) intervals (Figure 6B). The baseline-corrected results presented in Figure 3F show that the time period by region interaction was also significant for high gamma power, with the difference in high gamma power between the two regions increasing after stimulus onset.

### Alpha-power suppression in anterolateral Heschl’s gyrus reflects changes in both spectral slope and narrowband alpha power

Finally, we wanted to establish whether the suppressed low frequency activity in alHG reflected a reduction in the strength of a narrowband oscillation that is present before stimulus onset, or could be better characterized as a change in the exponent *χ* (“slope”) of the scale-free 1/*f*^χ^ component of the power spectrum. The latter may reflect properties of neural circuits distinct from those that generate narrowband oscillations (He, 2014; Gao, 2015; Podvalny et al., 2015; Voytek et al., 2015; Becker et al., 2018). The analyses described so far, in which power at each frequency was normalized by its value during a pre-stimulus baseline, cannot differentiate between these two contributions to neural power spectra.

To address this issue, for each hemisphere and region (pmHG, alHG; power averaged across contacts), we separately estimated the linear slope of the spectrum (on a log-log scale) and the residual narrowband oscillatory power after whitening by removal of the 1/*f*^χ^ component (i.e., the slope). The slope of the spectrum was shallower during sound presentation compared to the pre-stimulus period in alHG (*p* = 0.039, *r*_*e*_ = 0.692), but not in pmHG (*p* = 0.742, *r*_*e*_ = 0.128; interaction: *p* = 0.008, *r*_*e*_ = 0.812; Figures 7A), accounting for some proportion of the low-frequency power changes described earlier. Critically, an additional narrowband oscillatory peak between 7 and 10 Hz was present in the whitened power spectrum (i.e., after slope removal) during the pre-stimulus period (Figure 7B). Power in the alpha (7-10 Hz) frequency band was larger in the pre-stimulus-onset time window (–0.6 s to –0.1 s) compared to power during sound presentation (0.5 s to 1.2 s) in pmHG (*p* = 0.023, *r*_*e*_ = 0.768) and alHG (*p* = 0.016, *r*_*e*_ = 0.768), and this effect was larger in alHG compared to pmHG (interaction: *p* = 0.023, *r*_*e*_ = 0.737; Figure 7B).

**Figure 7.**
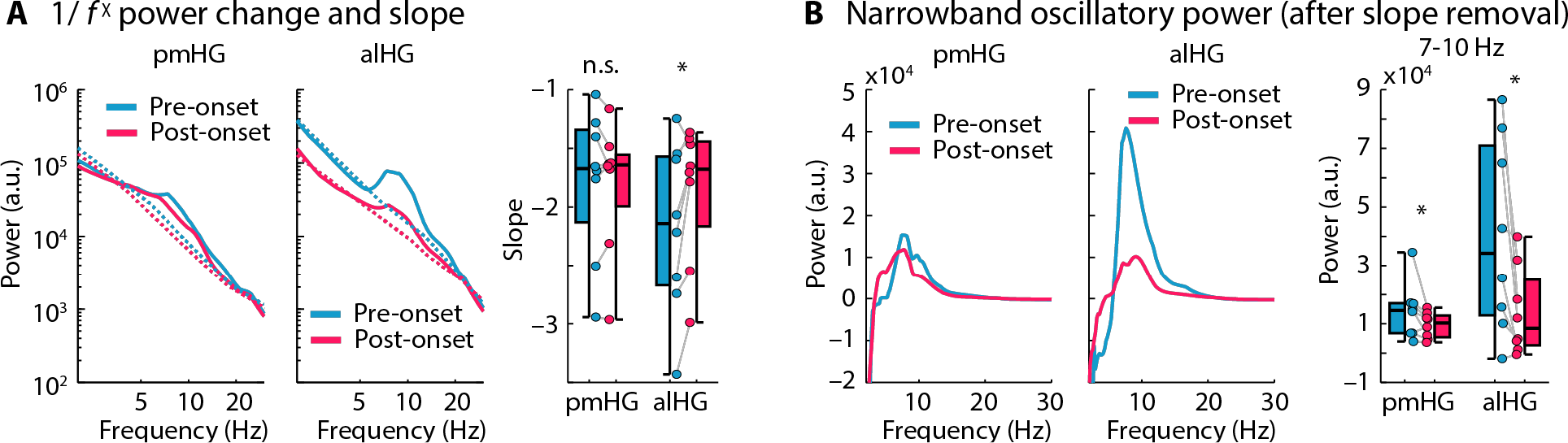
Separation of 1/*f*^χ^ slope and narrowband oscillatory power. **A:** Low frequency (2-30 Hz) power spectrum for pmHG and alHG during the pre-stimulus-onset (–0.6 s to –0.1 s) and post-stimulus-onset (0.5 s to 1.2 s) time windows, plotted on a log-log scale. The dotted lines reflect the best linear fit to the log-transformed data (individual fits averaged across hemispheres). Box plots and individual data points show the slope of the best linear fit in the pre-stimulus and post-stimulus time windows for the pmHG and alHG. **B:** Low frequency (2-30 Hz) power spectrum after removal of the 1/*f*^χ^ component for the pmHG and alHG during the pre-stimulus and post-stimulus time windows, plotted on a linear scale. Box plots and individual data points show the power in the 7-10 Hz band in the pre-stimulus and post-stimulus time windows for pmHG and alHG after removal of the 1/*f*^χ^ component. pmHG - posteromedial Heschl’s gyrus; alHG - anterolateral Heschl’s gyrus. ^*^*p* ≤ 0.05, n.s. - not significant.

## Discussion

Intracranial recordings revealed distinct profiles of spectral power across anatomically and functionally defined auditory cortical fields, both prior to, and during, auditory stimulation. Stimulus-related activity in the high gamma range was greater in posteromedial Heschl’s gyrus (pmHG, a primary field) than in anterolateral Heschl’s gyrus (alHG, a non-primary field; Brugge et al., 2009; Steinschneider et al., 2014). Lower frequency oscillations also differed across these areas, most markedly in the 7-10 Hz (“alpha”) range. In the absence of auditory stimulation (i.e. prior to stimulus onset), alpha power was stronger in alHG than in pmHG; this difference began to diminish following the onset of speech as alHG alpha power decreased. The suppression persisted until auditory stimulation ended, and could not be accounted for by a change in the exponent *χ* (“slope”) of the 1/*f*^χ^ component of the spectrum alone (Gao, 2015; Podvalny et al., 2015).

It has been proposed that one function of alpha oscillations throughout the brain is to reduce cortical excitability, limiting spontaneous or task-irrelevant processing (Klimesch et al., 2007; Jensen and Mazaheri, 2010). One might expect such a diminution in activity to manifest as reduced high gamma power, which is considered a proxy for asynchronous neural spiking (Crone et al., 2001; Mukamel et al., 2005; Manning et al., 2009). The anatomical dissociation we observed between alpha-power suppression (more anterolateral) and high gamma-power increases (more posteromedial) implies that any such antagonistic relationship does not hold as a fixed ratio across all sites. One interpretation is that primary cortex is ready to process sensory input somewhat indiscriminately, reflected in low spontaneous alpha activity, whereas non-primary areas are selectively engaged through suppression of ongoing alpha oscillations following the onset of particular stimuli (de Pesters et al., 2016). We observed such a stimulus dependency: alpha-power suppression for clear speech began earlier and was more pronounced than for spectrally degraded speech. Here we have shown that this effect, previously reported in scalp EEG (Obleser and Weisz, 2012), holds in non-primary auditory cortex.

Alpha oscillations have also been linked with predictive coding accounts of brain function (Friston and Kiebel, 2009). Activity in low frequency bands is thought to carry predictions of the causes of sensory data from higher to lower hierarchical levels (von Stein et al., 2000; Bastos et al., 2012; Fontolan et al., 2014; Chao et al., 2018), or to reflect the precision of such predictions (Bauer et al., 2014; Sedley et al., 2016; Auksztulewicz et al., 2017). The finding that alpha oscillations propagate from non-primary to primary visual cortex in monkeys (van Kerkoerle et al., 2014) is consistent with a feedback role. It has also been argued that alpha oscillations dominate in infragranular cortical layers that contain dense descending projections, and that this further implicates them in feedback processing (Maier et al., 2010; Bollimunta et al., 2011; Buffalo et al., 2011; Smith et al., 2013; Bastos et al., 2018; Bonaiuto et al., 2018). However, different referencing and analysis approaches have revealed alpha sources in all cortical layers (Haegens et al., 2015; Halgren et al., 2018). In any case, in order to adequately describe electrophysiological activity during auditory processing, predictive coding models will need to account for the differences we report between auditory fields both in baseline alpha power and in its stimulus-related suppression. Simultaneous laminar recordings from core and belt areas in non-human primates would be beneficial in isolating input and output layers at different hierarchical levels.

Linking the observed spectral signatures of primary and non-primary cortex to underlying cytoarchitectural, myeloarchitectural, and chemoarchitectural properties is complicated by the lack of a single agreed parcellation of the human superior temporal plane, and by the variability of Heschl’s gyrus morphology across individuals (von Economo and Horn, 1930). Our pmHG sites probably belong to core regions termed Te1.1/Te1.0 (Morosan et al., 2001) or AI (Rivier and Clarke, 1997; Wallace et al., 2002) and the alHG sites to Te1.2 (Morosan et al., 2001) or ALA (Wallace et al., 2002). Regardless of nomenclature, the former likely had a more developed granular layer with denser myelinated input from thalamus, and the latter more extensive long-range connections to non-sensory cortices (Hackett et al., 2001, 2014; Munoz-Lopez et al., 2010). These properties may give rise to preferred timescales of activity in primary and non-primary cortices corresponding to different spectral profiles (Honey et al., 2013; Murray et al., 2014). In terms of chemoarchitecture, the distribution of receptors and enzymes underlying cholinergic transmission in particular differs across the two fields (Hutsler and Gazzaniga, 1996; Zilles et al., 2002). This is relevant as the cholinergic system influences cellular excitability (Cox et al., 1994; Hsieh et al., 2000) as well as the stimulus coding capacity of a neural population (Minces et al., 2017; Schmitz and Duncan, 2018), and disrupting it pharmacologically can reduce alpha power (Osipova et al., 2003; Bauer et al., 2012; Eckart et al., 2016).

Alpha-power suppression was observed not only in anterolateral Heschl’s gyrus. Proximal contacts on the planum temporale, planum polare, and superior temporal gyrus showed similar activity patterns, with alpha suppression decreasing with distance from the centre of anterolateral Heschl’s gyrus. Whether these measurements reflect (suppression of) a single volume-conducted source of alpha activity or separate oscillators has yet to be established. A related observation is that the spectral profiles in posterolateral superior temporal gyrus and pmHG are more similar to each other than are profiles between the two subdivisions of Heschl’s gyrus. This is consistent with the identification of a posterolateral superior temporal (PLST) auditory area with similar functional properties to and direct connections with core auditory cortex (Howard et al., 2000; Brugge et al., 2003; Nourski et al., 2013, 2014).

The dominance of posterior alpha sources in recordings from the scalp can hinder detection of distinct alpha activity from the temporal lobe. Furthermore, alpha-power changes following the presentation of brief auditory stimuli, such as clicks, tones or syllables, are likely dominated by the broadband low-frequency component of the evoked response (also visible in Figure 3A and C). Such stimuli may be too short to elicit the sustained suppression we observed, which reached a maximum around 500-ms after stimulus onset and continued for the duration of the sentence. Nonetheless, magnetoencephalography, which is more selectively sensitive to tangentially-oriented dipoles that dominate on the superior temporal plane than EEG (Hämäläinen et al., 1993), has revealed likely temporal-lobe alpha sources without establishing their precise location (Tiihonen et al., 1991; Lehtelä et al., 1997; Weisz et al., 2011; Leske et al., 2015; Magazzini et al., 2016). Depth electrode recordings from Heschl’s gyrus in humans are relatively rare, and analysis of such data has largely focused on the evoked LFP or on high gamma activity (Nourski and Howard, 2015). Relevant exceptions include a multi-center study of resting state oscillatory activity (Frauscher et al., 2018). In that study alpha band peaks were present in activity from superior temporal gyrus but not Heschl’s gyrus, nor elsewhere on the posterior superior temporal plane. These findings are somewhat consistent with our pre-stimulus results but they included little if any data from anterolateral Heschl’s gyrus. Another study, in which posteromedial Heschl’s gyrus was sparsely sampled, did not consistently reveal changes in alpha activity in response to sound, although broadband low-frequency activity in one patient was suppressed in auditory association cortex on the lateral surface (Fontolan et al., 2014). Finally, Podvalny et al. (2015) noted stimulus-related alpha suppression over and above a 1/*f*^χ^ slope change in intracranially-implanted auditory areas, but did not report anatomical details. One contribution of the current report is to functionally identify sites as primary versus non-primary auditory cortex in all subjects, based on known neurophysiological response properties.

For our primary analyses we compared power pre-stimulus to that more than 500-ms after sentence onset, excluding transient evoked activity. We did not consider oscillatory phase, which is known to be important for perception (Busch et al., 2009; van Rullen, 2016) and may underpin communication between different brain regions (Palva and Palva, 2011). Indeed, a reduction in alpha power such as we observed in non-primary cortex may occur alongside increased inter-areal alpha-band phase synchrony that has been argued to support cognitive function such as working memory (Freunberger et al., 2008, 2009; Doesburg et al., 2009). A full account linking alpha activity in different cortical fields to auditory processing will need to take both power and phase into consideration.

In summary, we report strong evidence for a source of alpha oscillations in alHG that is suppressed during prolonged auditory stimulation. This suppression is more pronounced when subjects listen to clear than to degraded speech, and drops off with distance from alHG throughout superior temporal cortex. The suppression cannot be explained solely by a change in the exponent χ (“slope”) of the 1/*f*^χ^ component of the spectrum, and has a different anatomical distribution to induced high gamma activity, which is strongest in pmHG. Theories concerning the generation and function of alpha oscillations should account for their differential manifestation and stimulus-sensitivity in primary and non-primary auditory cortex.

